# Prevalence and genotyping of *Trichomonas gallinae* in Riyadh, Saudi Arabia

**DOI:** 10.1101/675033

**Authors:** Mohammed Fahad Albeshr, Abdulwahed Fahad Alrefaei

## Abstract

*Trichomonas gallinae* is a single-celled protozoan parasite and causative agent of trichomonosis. Trichomonosis is a disease with a worldwide distribution, and has recently been highlighted as a pandemic threat to several wild bird species. The aim of this study is to investigate the prevalence and genotypic diversity of *Trichomonas gallinae* in Riyadh, Saudi Arabia. A total of 273 oral swab samples from birds were collected and tested for *T. gallinae* infection, and the overall prevalence of *Trichomonas gallinae* in these samples was 26.4% (72 of 273). We compared the rates of infections with *T. gallinae* between wild and domestic pigeons and found that the infection was significantly higher in domestic pigeons; a finding which should be considered by the Ministry of Environment, Water and Agriculture of Saudi Arabia when evaluating the role of poultry markets in the transmission of trichomonosis. Sequence analyses of the internal transcribed spacer (ITS) region indicated genetic variation in these parasite strains, as found in the samples collected. Among 48 sequences 15 different ribotypes were found, 12 of which were novel, and 3 of which were previously-described ribotypes: ribotype A, C, and II. This study demonstrates the diversity of *T. gallinae* strains in Saudi Arabian birds for the first time, and revealed that ribotypes A and C are predominant among Riyadh birds.

## 1. Introduction

Parasitic diseases can negatively influence human health and domestic animal fitness (Cox-Witton et al., 2014). They can also increase the extinction risk of wild and captive natural populations by reducing the survival rate and reproductive fitness (Lachish et al., 2007, Höner et al., 2012). *T. gallinae* was first described by Rivolta in 1878 when it was found in an oral swab from a pigeon. Trichomonosis is a serious disease that caused by the single celled flagellate protozoan parasite *T. gallinae* (Stabler, 1947b), and is a global infectious disease usually affecting Columbiformes as primary hosts (Bunbury, 2011, Stockdale et al., 2015, Abouhosseini Tabari and Youssefi, 2018, Swinnerton et al., 2005). However, it can jump from Columbiformes to many different bird species around the world, including Psittacines (McKeon et al., 1997), Galliformes (Mantini et al., 2009), Strigiformes (Rogers et al., 2016, Cousquer, 2005), raptors (Falconiformes) (Sansano-Maestre et al., 2009), and Passerines (Burton and Doblar, 2004, Lehikoinen et al., 2013)

The poultry market in the capital of Saudi Arabia contains a wide variety of birds of many different species. Some of these birds, such as chickens, ducks, and pigeons, derive from local poultry farms; however, some, such as ornamental birds, parrots, and wild migrating birds such as falcons, are importated from outside the country. Such birds are likely to have been exposed to a variety of parasitic diseases including trichomonosis, and thus pose a spill-over threat to sympatric wild populations. This disease can also be transmitted to local poultry farms and cause significant economic losses (Reid et al., 1964).

The infection can be transmitted directly via squab feeding, when adult males and females exchange food as breeding behavior, or indirectly via sharing food or water (Bunbury et al., 2007, Swinnerton et al., 2005). Trichomonosis can be directly diagnosed through symptoms such as lesions in the digestive or in the upper respiratory tracts and swelling in the throat, nostrils, and eyes (Stabler, 1954, Narcisi et al., 1991). Death occurs due to breathing difficulties and/or starvation (Stabler, 1947a, Stabler, 1957, Tasca and De Carli, 2003). However, infected adult birds may not show signs of the disease, since they can develop a tolerance to the infection (Bunbury et al., 2007).

Trichomonosis has threatened several rare wild bird species around the world. For example, it was introduced to Mauritius with exotic bird species and transmitted to the endemic bird species on the island, including the echo parakeet and pink pigeon (Bunbury et al., 2007, Swinnerton et al., 2005). As a result, the populations of these two avian species have drastically declined in number in Mauritius (Bunbury et al., 2007, Swinnerton et al., 2005). In addition to wildlife concerns, there are increasing concerns that this disease could negatively affect the economic sectors and recreational values of certain countries (Gortázar et al., 2007).

Several strains of *T. gallinae* with differing virulence have been identified. For example, Sansano-Maestre et al. (2009) used the 5.8S ribosomal RNA (renal) region and restriction fragment length polymorphism (RFLP) markers to identify the strains of *T. gallinae* in wild or domestic pigeons (*Columba livia*) and birds of prey in eastern Spain. They found two genotypes in isolates from the Columbiformes and raptors. In these isolates, the genotypes isolated from the Columbiformes were more prevalent in the Columbiformes, and the genotypes isolated from the raptors were more prevalent in the raptors, displaying lesions. In addition, when comparing the sequencing of the 5.8S rDNA region and internal transcribed spacer (ITS) region in *T. gallinae* in Mauritian birds, including the turtle-dove (*Streptopelia picturata*) and the pink pigeon (*Columba mayeri*) (Gaspar da Silva et al., 2007), there were no variations between the isolates from these species.

In Saudi Arabia, investigations of *T. gallinae* in endemic and commensal birds are rare, and most research has been done in the family Falconidae (Samour and Naldo, 2003, Naldo and Samour, 2004). Here, we investigated the prevalence and genotypes of *Trichomonas gallinae* infections in avian species at the bird auction in Riyadh, comparing the prevalence to that of wild bird samples obtained locally. We provide evidence that the bird auction in Riyadh may be a source of infection in the area. A strength of this study is its focus on multiple species in limited geographic area, a clear advantage to similar studies performed in this region of the world previously.

## 2. Material and Methods

We examined a total of 273 oral swabs from 5 bird species, collected between 2018 and 2019 in Riyadh, Saudi Arabia (Table 1). Of these birds, 90 individuals (73 Common mynah and 17 feral pigeons) were caught in the wild using mist-net captures between March and May 2018.

**Table 1.**
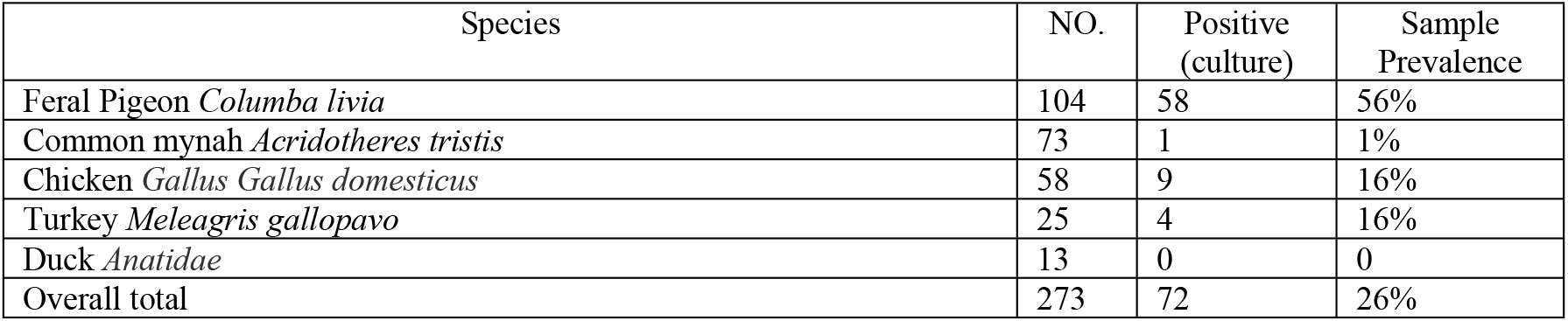
Prevalence (%) of *Trichomonas gallinae* infection in 5 bird species, sampled in Riyadh, Saudi Arabia.

| Species | NO. | Positive (culture) | Sample Prevalence |
| --- | --- | --- | --- |
| Feral Pigeon *Columba livia* | 104 | 58 | 56% |
| Common mynah *Acridotheres tristis* | 73 | 1 | 1% |
| Chicken *Gallus Gallus domesticus* | 58 | 9 | 16% |
| Turkey *Meleagris gallopavo* | 25 | 4 | 16% |
| Duck *Anatidae* | 13 | 0 | 0 |
| Overall total | 273 | 72 | 26% |

Briefly, the oral swab samples were obtained from the oral cavity to the crop with sterile cotton swabs moistened with distilled water. The swab was gently moved four or five times in a figure-eight motion around the oral cavity and crop mucosa to ensure that a sufficient sample of mucosal and trichomonad cells was obtained. The swabs were then immediately inoculated individually into InPouchTM TV culture kits (BioMed Diagnostics) using the manufacturer’s guidance and instructions. The samples were transferred to the laboratory and kept at 36°C in a sanitized incubator. All culture samples were examined 24 hours post-incubation under a microscope, and thereafter at 24-hour intervals for up to seven days to check for *T. gallinae* parasite growth. Positive samples were stored in 5% DMSO at −80 °C for future work. During swabbing, the birds were inspected in the oral cavity for signs of infection with *T. gallinae*, including lesions around the oral cavity and eyes. However, a diagnosis of infection with *T. gallinae* was only confirmed when physical symptoms were validated by microscopic examination.

DNA extraction from positive samples was performed using DNAzol® following the manufacturer’s instructions. The InPouchTM TV cultures containing *Trichomonas gallinae* were transferred to 1000 μl Eppendorf tubes and then centrifuged at 9,000 rpm for 3 min. The supernatant was removed whereas the pellets were retained, and 500 μL DNAzol was added to the samples and pipetted up and down to lyse the cells, and the samples were subsequently centrifuged at 5,000 rpm for 4 min at 4 °C. To precipitate DNA, the resulting of supernatant was transferred to a new Eppendorf tube. 500 μL of 100% ethanol was added to each tube, followed by mixing through inversion and 3 min of centrifugation at 4,000 rpm in 4 °C. The liquid was then removed and the DNA pellet was washed in 75% ethanol. The DNA pellets were left to air-dry for 4 min before adding 100 μL of nuclease-free water to re-suspend the DNA. The extracted of DNA was left in −20 °C for subsequent work. The quantity of DNA for each isolate was estimated using spectrophotometry, based on an absorbance reading of 260 nm. The extracted DNA was confirmed visually on a 1% of agarose gel stained in ethidium bromide, under an ultraviolet transilluminator.

PCR was conducted using the method described by Robinson et al. (2010) to amplify the ITS region, using the forward primer TFR1 (5’TGCTTCAGTTCAGCGGGTCTTCC3’) and the reverse primer TFR2 (5’CGGTAGGTGAACCTGCCGTTGG-3’) (Felleisen, 1997; Gaspar da Silva et al., 2007). PCR was carried out in 25 μL of mixture volume per sample, containing 15 μL Green Master Mix (2X, USA), 3 μL of forward and reverse primer (Eurofins Genomics, Germany), 3 μL of ddH2O, and 1 μL of template DNA. Each PCR was run with a negative control without any template DNA, as well as a positive control containing *T. gallinae* DNA.

The PCR reactions were performed using the following temperature cycle: initial denaturation at 94 °C for 15 min, followed by 35 cycles with denaturation at 94 °C for 1 min, annealing at 65 °C for 30 seconds, extension at 72 °C for 1 min, and a final extension at 72 °C for 5 min.

We confirmed PCR amplification under ultraviolet light used a 1% agarose gel stained in ethidium bromide. A band of about 350 base pairs (bp) in size was confirmed using a Ready-LoadTM 100 bp DNA ladder (Promega, USA). We submitted the positive PCR products for sequencing to Macrogen Inc (Seoul, Korea).

The molecular and phylogenetic relationship of sequences of trichomonad parasites were determined using MEGA software version 7 (Tamura et al., 2011) and CLUSTALX 2.1 (Thompson et al., 1997). Published *Trichomonas gallinae* sequences found in the NCBI GenBank database were used to compare the ITS regions (S1 Table). Phylogenetic trees of the datasets obtained from the ITS1/5.8S and rRNA/ITS2 regions were constructed separately using the NJ and the Tamura-Nei models. These were used to analyze relationship between taxa through nucleotide sequence analysis (Saitou and Nei, 1987, Tamura et al., 2011). We used the Felsenstein’s bootstrap testing to calculate the associate taxa grouped in the bootstrap values (1,000 times) (Felsenstein, 1985).

## 3. Results

Of the 273 birds examined in this study, 72 (26.4%) tested positive for *Trichomonas gallinae* infection, including 56% of feral pigeons (FP), 1% of common mynah (CM), 16% of chickens (CG), 16% of turkeys (TM) and 0% of ducks (DA) sampled (Table 1; Figures 1 and 2). There were significant differences in infection prevalence between species, (χ^2^ = 79.346, d.f. = 4, p < 0.001).

**Figure 1.**
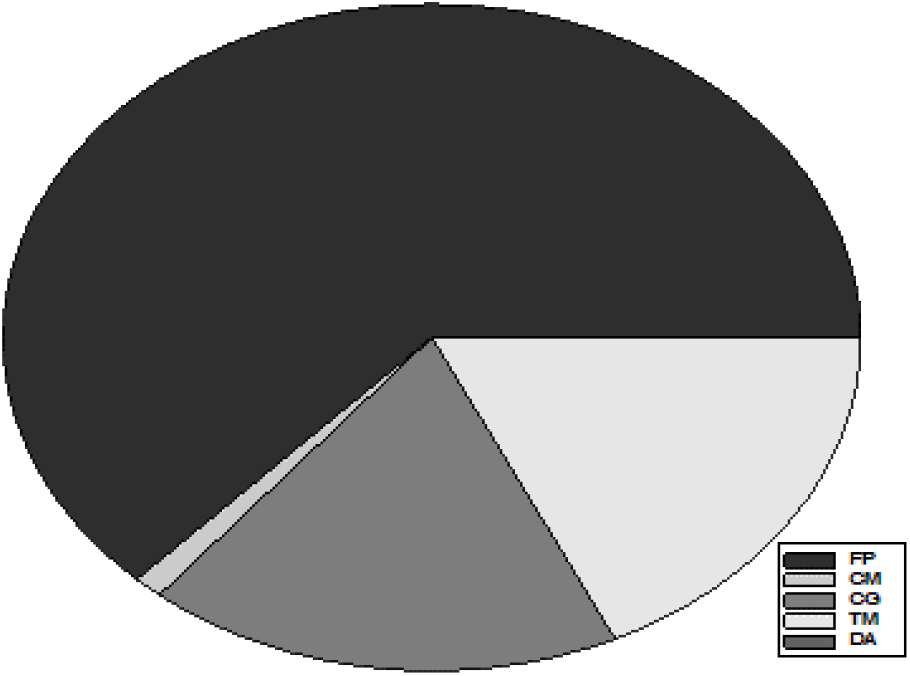
Proportion of prevalence (%) of *Tichomonas gallinae* infection across five bird species (FP = Feral Pigeon, CM = Common mynah, CG = Chicken *Gallus Gallus domesticus*, TM = Turkey *Meleagris gallopavo* and DA = Duck *Anatidae*).

**Figure 2.**
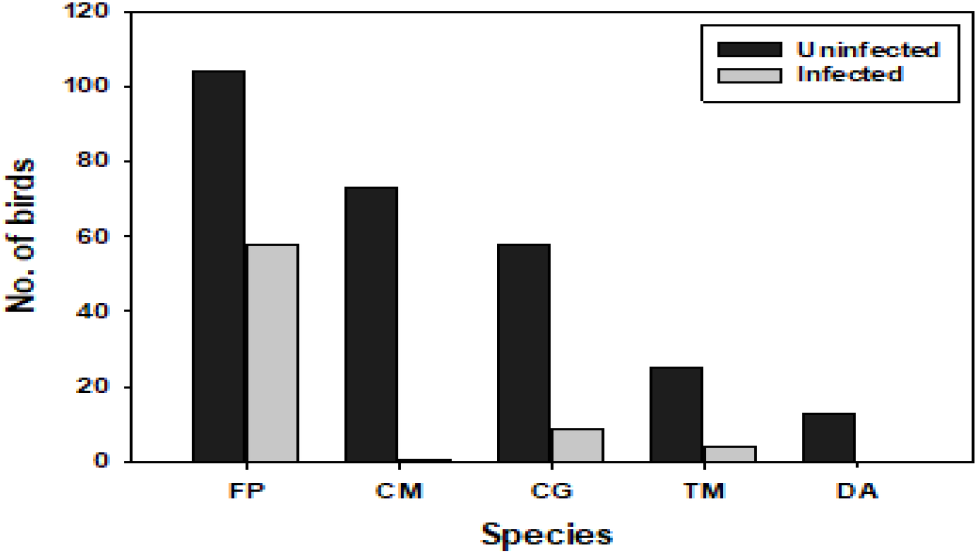
Number of infected and uninfected birds with *T. gallinae* in five bird species (FP = Feral Pigeon, CM = Common mynah, CG = Chicken *Gallus Gallus domesticus*, TM = Turkey *Meleagris gallopavo* and DA = Duck *Anatidae*).

**Table 2.** Overview of isolate ID, birds species, clinical signs, characteristic lesions, and culture using the ITS ribotype of *Trichomonas gallinae* in birds found in Riyadh, Saudi Arabia.

|  | Case ID | Host species | Bird status | Clinical signs | Lesions | Culture extract | ITS ribotype |
| --- | --- | --- | --- | --- | --- | --- | --- |
| 1 | R1 | Feral pigeon | Alive |  | - | + | C |
| 2 | R3 | Feral pigeon | Alive | + | + | + | C |
| 3 | R4 | Feral pigeon | Alive | - | - | + | A |
| 4 | R5 | Feral pigeon | Alive |  | - | + | C |
| 5 | R6 | Feral pigeon | Alive | - | + | + | A |
| 6 | R7 | Feral pigeon | Alive | - | - | + | KSA11 |
| 7 | R8 | Feral pigeon | Alive | - | + | + | KSA10 |
| 8 | R9 | Feral pigeon | Alive | - | - | + | KSA7 |
| 9 | R10 | Feral pigeon | Alive | + | + | + | C |
| 10 | R12 | Feral pigeon | Alive | - | + | + | KSA1 |
| 11 | R13 | Feral pigeon | Alive | - | - | + | C |
| 12 | R16 | Feral pigeon | Alive | + | + | + | C |
| 13 | R17 | Feral pigeon | Alive | - | - | + | C |
| 14 | R18 | Feral pigeon | Alive | - | - | + | C |
| 15 | R19 | Feral pigeon | Alive | - | + | + | II |
| 16 | R24 | Feral pigeon | Alive | - | - | + | C |
| 17 | R25 | Feral pigeon | Alive | - | + | + | KSA4 |
| 18 | R27 | Feral pigeon | Alive | - | + | + | KSA9 |
| 19 | R28 | Feral pigeon | Alive | - | + | + | KSA8 |
| 20 | R29 | Feral pigeon | Alive | - | + | + | KSA6 |
| 21 | R30 | Feral pigeon | Alive | - | + | + | C |
| 22 | R31 | Feral pigeon | Alive | - | + | + | KSA2 |
| 23 | R32 | Feral pigeon | Alive | - | - | + | A |
| 24 | R34 | Feral pigeon | Alive | - | - | + | C |
| 25 | R35 | Feral pigeon | Alive | - | - | + | KSA3 |
| 26 | R37 | Feral pigeon | Alive | + | + | + | C |
| 27 | R38 | Feral pigeon | Alive | - | - | + | A |
| 28 | R41 | Feral pigeon | Alive | + | + | + | A |
| 29 | R47 | Chicken | Alive | - | - | + | A |
| 30 | R49 | Chicken | Alive | + | + | + | A |
| 31 | R50 | Chicken | Alive | - | - | + | A |
| 32 | R51 | Chicken | Alive | - | - | + | A |
| 33 | R52 | Chicken | Alive | - | - | + | A |
| 34 | R53 | Chicken | Alive | - | - | + | A |
| 35 | R55 | Feral pigeon | Alive | - | - | + | C |
| 36 | R57 | Turkey | Alive | - | - | + | A |
| 37 | R58 | common mynah | Alive | - | - | + | A |
| 38 | R59 | Turkey | Alive | - | - | + | A |
| 39 | R60 | Turkey | Alive | - | - | + | A |
| 40 | R61 | Turkey | Alive | - | - | + | A |
| 41 | R62 | Feral pigeon | Alive | - | - | + | C |
| 42 | R65 | Feral pigeon | Alive | + | + | + | C |
| 43 | R67 | Feral pigeon | Alive | + | + | + | KSA5 |
| 44 | R68 | Feral pigeon | Alive | + | + | + | KSA12 |
| 45 | R69 | Feral pigeon | Alive | + | + | + | C |
| 46 | R70 | Feral pigeon | Alive | + | + | + | C |
| 47 | R71 | Feral pigeon | Alive | + | + | + | A |
| 48 | R72 | Feral pigeon | Alive | + | + | + | C |

**Fig. 1.**
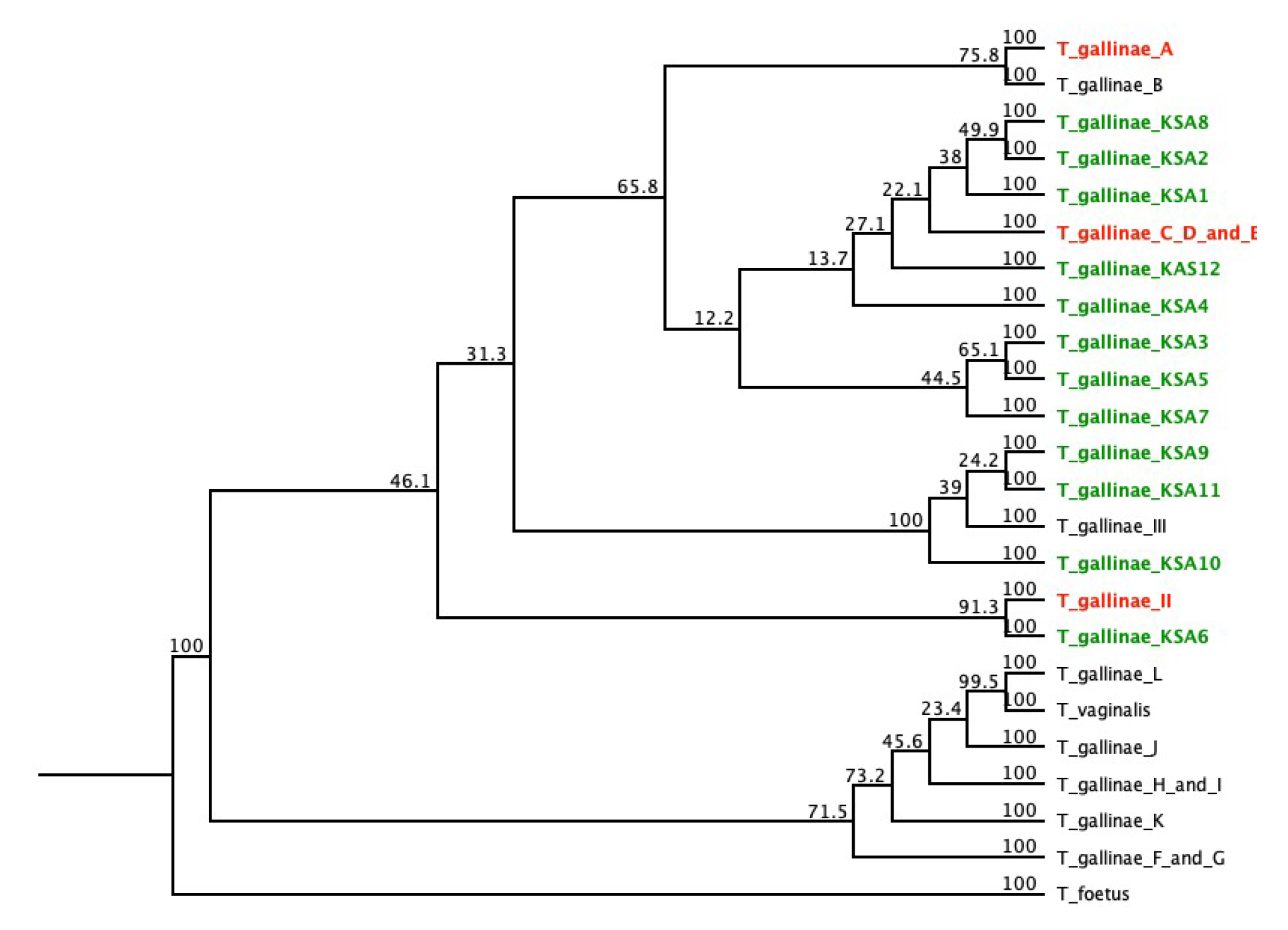
The phylogenetic tree shows the relationship of *T. gallinae* using the NJ method for the ITS region from sequences described in Table 2. *Tritrichomonas foetus* was included in this study as an outgroup. The novel ribotypes identified in this study are marked in green. Reference sequences are labeled with their lineage in black, and the red-labelled sequences indicate previously identified ITS ribotypes of the samples analyzed in this study.

The proportions of infections differed significantly between wild pigeons (n = 17), and poultry market’s pigeons (n = 87), z= 2.129; *p* = 0.033. Lesions associated with trichomoniasis were examined in the birds, revealing that 51.73% of infected pigeons and 33.34% of infected chickens showed lesions inside and outside of the oral cavity and eyes.

In the 72 samples, the ITS region was amplified through PCR using ITS-specific primers, and a fragment of about 300 bp was obtained (Table 1). Of these positive PCR samples, 48 were randomly selected and sequenced (Table 2). The phylogenetic tree for the ITS region contained 12 different novel lineages, including three previously described ribotypes: ribotypes A (29 samples), C (6), and II (one sample) (Figure 1). Most samples clustered in lineage A (Lawson et al., 2011b) (GenBank: GQ150752), which was found in 17 samples. This included columbids (feral pigeon), passerines (common mynah), and galliform (chicken, turkey). 18 samples of columbids (feral pigeon) were predominantly infected by lineage C (Gerhold et al., 2008) (GenBank EU215362). One isolate of *Trichomonas gallinae* obtained from feral pigeons that was grouped in the same branch as a sequence of the *Trichomonas gallinae* ITS region lineage II, obtained from a racing pigeon (Grabensteiner et al., 2010) (GenBank FN433474). Interestingly, comparison of the complete ITS region of all novel determined sequences in this study revealed 12 different sequence lineages (KSA1–KSA12). These demonstrated a degree of sequence divergence between each other, and each of them formed a separate branch. All of these occurred in columbids (feral pigeon). The sequences types (ITS; KSA1–KSA12) were novel, reported for the first time in the present investigation in Riyadh, Saudi Arabia.

## 4. Discussion

Trichomonosis is an avian disease that is caused by *T. gallinae*, and is a widespread ailment affecting different bird species worldwide, characterized by a great variation in strain virulence and pathogenicity. In the last few years, interest surrounding investigating *T. gallinae* strains has increased and spurred the introduction of new molecular techniques. In this paper, we studied the prevalence and genotypes of *Trichomonas gallinae* infections in avian species at the bird auction in Riyadh, including birds that were caught in the wild. We demonstrate for the first time the strains of *T. gallinae,* using the ITS ribotype to compare the genetic variation of *T. gallinae* isolated from different bird species in Riyadh, Saudi Arabia.

We found *Trichomonas gallinae* to be present in four of the bird species examined in this study, confirming the cases in Saudi Arabia, with an overall prevalence rate of 26.4%. The prevalence of *Trichomonas gallinae* found in the poultry market in Riyadh was much higher in pigeons than other bird species. This was a predictable result, as feral pigeons have been considered the primary host of *T. gallinae* infections (Stabler, 1954, Stabler, 1951). However, this parasite can seemingly be transmitted to other species involved in this study, including the common mynah (Farooq et al., 2018), the *Gallus Gallus domesticus* chicken (Grabensteiner et al, 2010), the *Meleagris gallopavo* turkey, and the *Anatidae* duck (Grunenwald et al, 2018), Furthermore, prevalence rates were found to be significantly higher in market pigeons compared to those caught in the wild. This provides evidence for implicating the auction as a mode of transmission of *T. gallinae* among and within bird species.

*T. gallinae* genotypes were divergent between different bird species, although our sample size was too small to be conclusive. The results revealed the existence of several *T. gallinae* strains circulating in Saudi Arabia avifauna. A phylogenetic analysis was used to identify fifteen unique sequencing in this investigation, and these clearly divide into different branches using the ITS ribotype (Figure 1). 12 of them were novel, and three have been previously described. In this study, we further present data on the genetic diversity of *T. gallinae* found in Saudi Arabian birds, withdifferent nomenclatures to differentiate between them (Table S1). Of the 15 genetic lineages found in this study, three genotypes have been found in previous studies: genotype A (Lawson et al., 2011a), genotype C (Gerhold et al., 2008) and genotype II (Grabensteiner et al., 2010). Regarding the previously described lineages and NCBI, the lineages KSA1–KSA2 might be newly detected lineages, as they have not been described in previous studies. Furthermore, these lineages appear to be distinct from lineages A/B and C/D/E (Lawson et al., 2011a; Gerhold et al., 2008); thus, they may not be as common or widespread as lineages A, B, C, and II (Lawson et al., 2011a, Gerhold et al., 2008, Zimre-Grabensteiner et al., 2011). This is because they were found only in pigeons in Saudi Arabia. However, judging by the overall occurrence of these three genotypes, our findings propose a widespread distribution of those genotypes among different bird species (Gerhold et al., 2008; Lawson et al., 2011a; Chi et al., 2013; Farooq et al., 2018; Alrefaei et al., 2019).

In this study, we found that most birds were infected by *T. gallinae* ribotypes A and C. This is consistent with prior studies, which found the pathogenicity of ribotype A is strong compared to genotype C, as demonstrated by the dramatic decline in the UK finch population in 2007 (Robinson et al., 2010; Chi et al., 2013; Alrefaei et al., unpublished). The C genotype isolated in this study was identified in pigeons and chickens with lesions, and was an apparently pathogenic lineage. We also found this symptom in birds infected with a novel genotype, as described in this study. However, most of the birds infected with genotype A appeared to be healthy. Several pigeons and chickens with ulcerations in the oral cavity were also observed to be infected with genotype C or a novel genotype. In 2009, Sansano-Maestre et al. observed that for Spanish samples, genotype C tended to be less virulent than genotype A, commonly associated with macroscopic lesions. Since then, a virulent clonal isolate of genotype A has been associated with both pigeon and passerine pathology and mortality (Chi et al., 2013). However, it is likely that since virulence is a rapidly evolved trait (De Fine Licht HH, 2018), different strains within distinct genetic lineages will vary markedly in virulence. In support of this, our study seems to suggest that it is genotype C rather than a genotype A with which pathology is primarily associated. Further research globally on this avian parasitic species, its genetic diversity, and the association with pathogenicity will elucidate to what extent, if any, virulent traits can be ascribed to genetic lineages of this parasite.

## Acknowledgements

“The authors extend their appreciation to the Deanship of Scientific Research at King Saud University for funding this work through the Research Project No NFG-7-18-03-06, and we thank the Researchers Support Services Unit (RSSU) at King Saud University for their technical support”.

## Ethics statement

All procedures for the samples collection were carried out in strict accordance with the recommendations by the Research Ethics Committee (REC) at the King Saud University (Ethic Reference No: KSU-SE-19-77).

---

**Table S1.** List of *Trichomonas gallinae* ITS ribotypes used as reference strains in this study. *Trichomonas vaginalis* was included as an outgroup.

| Species (host) | Origin | ITS<br>ribotype | GenBank | GenBank |
| --- | --- | --- | --- | --- |
| *T. gallinae* (Greenfinch *Chloris chloris*) | UK | A<sup>a</sup> | GQ150752 | Lawson et al. (2011) |
| *T. gallinae* (Broad-winged hawk *Buteo platypterus*) | Florida, USA | B | EU215368 | Gerhold et al. (2008) |
| *T. gallinae* (Rock pigeon *Columba livia*) | Colorado, USA | C<sup>b</sup> | EU215362 | Gerhold et al. (2008) |
| *T. gallinae* (Common ground dove *Columbina passerina*) | Texas, USA | F | EU215358 | Gerhold et al. (2008) |
| *T. gallinae* (Common ground dove *Columbina passerina*) | Texas, USA | G | EU215359 | Gerhold et al. (2008) |
| *T. gallinae* (White-winged dove *Zenaida asiatica*) | Texas, USA | H | EU215360 | Gerhold et al. (2008) |
| *T. gallinae* (White-winged dove *Zenaida asiatica*) | Texas, USA | I | EU215361 | Gerhold et al. (2008) |
| *T. gallinae* (Mourning dove *Zenaida macroura*) | Texas, USA | J | EU215365 | Gerhold et al. (2008) |
| *T. gallinae* (Band-tailed pigeon *Patagioenas fasciata*) | Californi, USA | K | EU215367 | Gerhold et al. (2008) |
| *T. gallinae* (Cooper's hawk *Accipiter cooperii*) | Arizona, USA | L | EU215366 | Gerhold et al. (2008) |
| *T. gallinae* (racing pigeon *Columba livia*) | Austria | II | FN433474 | Grabensteiner et al. (2010) |
| *T. gallinae* (ferial pigeon *Columba livia*) | Riyadh, Saudi Arabia | KSA1 | MK771125 | current study |
| *T. gallinae* (ferial pigeon *Columba livia*) | Riyadh, Saudi Arabia | KSA2 | MK771126 | current study |
| *T. gallinae* (ferial pigeon *Columba livia*) | Riyadh, Saudi Arabia | KSA3 | MK77112<br>7 | current study |
| *T. gallinae* (ferial pigeon *Columba livia*) | Riyadh, Saudi Arabia | KSA4 | MK77112<br>8 | current study |
| *T. gallinae* (ferial pigeon *Columba livia*) | Riyadh, Saudi Arabia | KSA5 | MK77112<br>9 | current study |
| *T. gallinae* (ferial pigeon *Columba livia*) | Riyadh, Saudi Arabia | KSA6 | MK77113<br>0 | current study |
| *T. gallinae* (ferial pigeon *Columba livia*) | Riyadh, Saudi Arabia | KSA7 | MK77113<br>1 | current study |
| *T. gallinae* (ferial pigeon *Columba livia*) | Riyadh, Saudi Arabia | KSA8 | MK77113<br>2 | current study |
| *T. gallinae* (ferial pigeon *Columba livia*) | Riyadh, Saudi Arabia | KSA9 | MK77113<br>3 | current study |
| *T. gallinae* (ferial pigeon *Columba livia*) | Riyadh, Saudi Arabia | KSA10 | MK77113<br>4 | current study |
| *T. gallinae* (ferial pigeon *Columba livia*) | Riyadh, Saudi Arabia | KSA11 | MK77113<br>5 | current study |
| *T. gallinae* (ferial pigeon *Columba livia*) | Riyadh, Saudi Arabia | KSA12 | MK76502<br>9 | current study |
| *T. vaginalis* (Human *Homo sapiens*) | Brazil | NA | AY34918<br>4 | Kleina et al. 2004 |
| *T. foetus* | France | NA | DQ24391<br>1 | Duboucher et al.<br>(2006) |

